# Cocaine-Induced Gene Regulation in D1 and D2 Neuronal Ensembles of the Nucleus Accumbens Revealed by Single-Cell RNA Sequencing

**DOI:** 10.1101/2024.06.19.599762

**Authors:** Philipp Mews, Autumn VA Mason, Emily G Kirchner, Molly Estill, Eric J Nestler

## Abstract

Cocaine use disorder is characterized by persistent drug-seeking behavior and a high risk of relapse, driven by lasting molecular and circuit adaptations in the nucleus accumbens. To explore the transcriptomic changes underlying these alterations, we employed fluorescence-activated nucleus sorting coupled with single-nucleus RNA sequencing to analyze D1 and D2 medium spiny neurons in this brain region of male mice subjected to acute cocaine exposure or to prolonged withdrawal from repeated cocaine exposure without or with an acute cocaine rechallenge. This approach allowed us to precisely delineate and contrast transcriptionally distinct neuronal subpopulations─or ensembles – across various treatment conditions. We identified significant heterogeneity within both D1 and D2 MSNs, revealing distinct clusters with unique transcriptional profiles. Notably, we identified a discrete D1 MSN population characterized by the upregulation of immediate early genes, as well as another group of D1 MSNs linked to prolonged withdrawal, uncovering novel regulators of withdrawal-related transcriptome dynamics. Our findings provide a high-resolution transcriptomic map of D1 and D2 MSNs, illustrating the dynamic changes induced by cocaine exposure and withdrawal. These insights into the molecular mechanisms underlying cocaine use disorder highlight potential targets for therapeutic intervention aimed at preventing relapse.

## INTRODUCTION

Substance use disorders, including cocaine use disorder (CUD), are characterized by enduring alterations in neural circuits and molecular pathways that drive the persistent compulsion to seek and consume drugs. The chronic and relapsing nature of addiction underscores the complexity of its underlying molecular pathology and the urgent need for advanced therapeutic strategies that target these fundamental alterations. Despite the growing crisis of cocaine use disorder, marked by rising overdose fatalities and the lack of approved pharmacotherapies^1–3^, our understanding of its molecular mechanisms remains incomplete. Central to the pathophysiology of addiction disorders, including CUD, is the mesolimbic dopaminergic circuitry, particularly the nucleus accumbens (NAc)^4–11^. This forebrain region plays a pivotal role in reward processing, motivational salience, and reinforcement learning, making it a prime target for investigating the neurobiological mechanisms driving addiction.

Historically, the vast majority of neurons (>90%) in the NAc have been categorized into two primary subtypes based on their dopamine receptor expression: D1 dopamine receptor-expressing medium spiny neurons (D1 MSNs) and D2 dopamine receptor-expressing medium spiny neurons (D2 MSNs)^11,12^. D1 MSNs are generally associated with the promotion of reward-related behaviors, whereas D2 MSNs are linked to the suppression of such behaviors^11–17^. Cocaine acts on these neurons to induce synaptic plasticity, a fundamental mechanism contributing to the drug’s addictive properties^18^. Activation of D1 MSNs typically enhances the rewarding effects of cocaine, while activation of D2 MSNs can inhibit these effects, although more recent research has painted a more complex picture^11,19–22^.

Recent studies utilizing RNA sequencing (RNAseq) of purified populations of these neurons have shown that D1 MSNs and D2 MSNs exhibit distinct transcriptional patterns at baseline and in response to cocaine, which are critical for their differential roles in addiction and reward processing^6,23^. Specifically, RNAseq data have revealed that D1 MSNs show upregulation of genes associated with synaptic plasticity, neurotransmitter release, and signal transduction pathways following repeated cocaine administration, thereby reinforcing cocaine’s rewarding effects and promoting addictive behaviors^24–28^. In contrast, in D2 MSNs, cocaine exposure typically activates genes associated with neurotransmitter inhibition, stress response, and neuroadaptive changes, potentially contributing to their role in countering cocaine’s rewarding effects^23,29^. These findings highlight that the divergent effects of cocaine on D1 and D2 MSNs stem from distinct transcriptional programs in these cells, reflecting their distinct influences on behavior.

Recent advances in single-nucleus RNAseq (snRNAseq) have provided a more nuanced understanding of these neuronal populations. Studies have revealed significant heterogeneity within D1 MSNs and D2 MSNs, indicating that subpopulations with distinct transcriptional profiles respond differently to cocaine exposure^24–26,30^. For instance, small subpopulations of D1 MSNs exhibit unique immediate early gene expression patterns in response to acute cocaine exposure in drug-naïve mice, which may play crucial roles in synaptic modifications underlying addiction and relapse^24,30^. This transcriptional complexity suggests that the traditional binary classification of NAc MSNs oversimplifies their roles and interactions within the broader neural circuitry involved in addiction^25^. Understanding these diverse transcriptional responses is critical for developing targeted interventions to address the multifaceted nature of cocaine addiction and its long-term consequences.

While prior research utilizing snRNAseq on bulk tissue has provided initial insights into the transcriptional responses of NAc cell populations to acute cocaine exposure, less effort has been devoted to examining the effects of chronic cocaine and this approach lacks the resolution to fully discern nuanced differences within neuronal subtypes. Consequently, the lasting effects of chronic cocaine on the transcriptomes of D1 and D2 MSNs remain largely unexplored. Traditional bulk RNAseq approaches group all neural subtypes together, diluting the specific signals from distinct neuronal populations and masking critical subtleties in gene expression dynamics.

The present study significantly advances the field by employing fluorescence-activated nucleus sorting (FANS) coupled with snRNAseq to first purify and then specifically analyze the transcriptomes of D1 and D2 MSNs. This study represents a pioneering effort to characterize the distinct transcriptomic responses of D1 and D2 MSNs with high resolution after acute cocaine exposure, after prolonged withdrawal following chronic cocaine, and in response to cocaine re-challenge during withdrawal. We identified unique molecular pathways and regulatory networks activated by cocaine and withdrawal across discrete D1 and D2 MSN subpopulations, offering potential novel targets for advanced treatments for CUD and drug relapse. Together, these findings provide a comprehensive understanding of the transcriptome dynamics underlying the enduring effects of cocaine on NAc neuronal function.

## RESULTS

To investigate the transcriptomic changes induced by cocaine and prolonged abstinence across distinct subpopulations of MSNs in the NAc, male mice were subjected to a regimen of 10 daily IP injections of either saline or cocaine (20 mg/kg). Following this initial phase, the animals underwent a 30-day withdrawal period before they were re-exposed to either a saline or cocaine injection and analyzed 1 hr later (Figure 1A). Utilizing FANS coupled with snRNAseq, we isolated and analyzed specific subpopulations of D1 and D2 dopamine receptor-expressing neurons (Figure 1B). This approach allowed for precise delineation and comparison of these subpopulations across four distinct conditions: saline control, acute cocaine exposure, prolonged withdrawal from chronic cocaine, and cocaine rechallenge after withdrawal. This methodology enabled an in-depth exploration of the dynamic transcriptomic landscape within the NAc, elucidating the cellular mechanisms underlying both the acute and enduring responses of MSNs to cocaine exposure and withdrawal.

**Figure 1:**
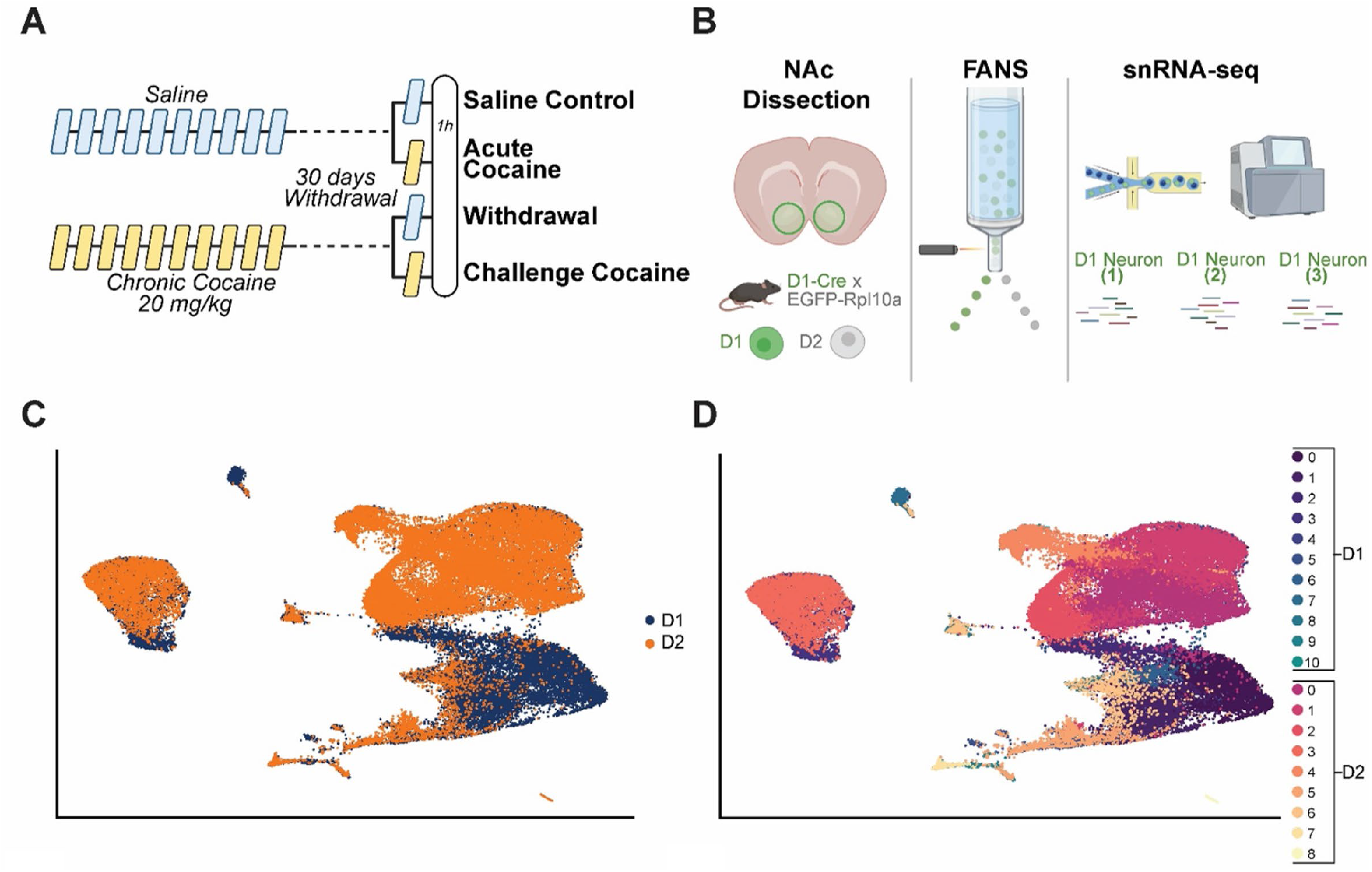
snRNAseq data analysis of D1 and D2 MSNs in the mouse NAc following cocaine exposure. C. UMAP visualization showing D1 and D2 MSN nuclei in a union UMAP. This plot provides an integrated view of the distribution of both neuron subpopulations. **D.** Union UMAP shows the different clusters for each subpopulation of D1 MSNs (D1 clusters 0-10) and D2 MSNs (D2 clusters 0-8).

Our analysis began with an overview of the snRNAseq data for all D1 and D2 dopamine receptor-expressing neurons. To explore the inherent heterogeneity of these cell populations, we applied the Seurat clustering algorithm, which utilizes a graph-based approach to cluster cells based on similar gene expression profiles. This method facilitated the identification of cell subpopulations according to their transcriptional signatures. The Uniform Manifold Approximation and Projection (UMAP) visualization provided an integrated perspective of the distribution of these D1 and D2 cell subpopulations, illustrating their spatial arrangement within a unified UMAP (Figure 1C). Virtually all of the isolated cells are MSNs based on known markers of this cell type, with only a very small (0.4%) subpopulation of D2 cells representing cholinergic interneurons (cluster D2_7), which are known to also express the D2 dopamine receptor and characterized by the expression of *Chat*, *Nxph1* and *Lhx8*^31–33^. These findings confirm prior research that the mouse lines used to demarcate D1 and D2 cells identify MSNs.

Our analysis identified 11 clusters within the D1 MSN population (clusters D1_0-10) and 9 clusters within the D2 MSN population (clusters D2_0-8), reflecting the substantial heterogeneity present within each of these cell populations (Figure 1D). We identified roughly 5% of D1 MSNs that express *Drd2* and 7% of D2 MSNs that express *Drd1*, confirming prior research that a small percentage of NAc MSNs express both receptors.

These UMAP plots underscore the distinct transcriptional identities of D1 and D2 MSNs, providing a foundational overview of the gene expression profiles within these neuronal subpopulations. This comprehensive clustering approach allowed for a systematic exploration of cellular diversity, enabling subsequent analyses to delve deeper into the transcriptional landscapes of individual clusters. Such detailed profiling is crucial for understanding the specific roles and responses of these neurons in the context of cocaine exposure.

### Transcriptomic Diversity of D1 MSNs

To explore the transcriptomic heterogeneity of D1 MSNs under the different treatment conditions, we first assessed the percentage makeup of each condition across the eleven identified D1 neuron clusters. The bar chart in Figure 2A illustrates the percentage composition of cells from each treatment condition (saline control, acute cocaine exposure, prolonged withdrawal, and cocaine rechallenge) within each cluster. This analysis reveals significant variations in the distribution of treatment conditions among the clusters, indicating how different treatments uniquely impact the transcriptional landscape of D1 MSNs. For instance, clusters such as D1_5 and D1_6 show a higher proportion of cells from the acute cocaine and cocaine challenge group, while clusters D1_7 and D1_8 are more populated by nuclei from the prolonged withdrawal group, suggesting that these clusters represent transcriptional states that play distinct roles in the immediate and long-term responses to cocaine (Figure 2A).

**Figure 2:**
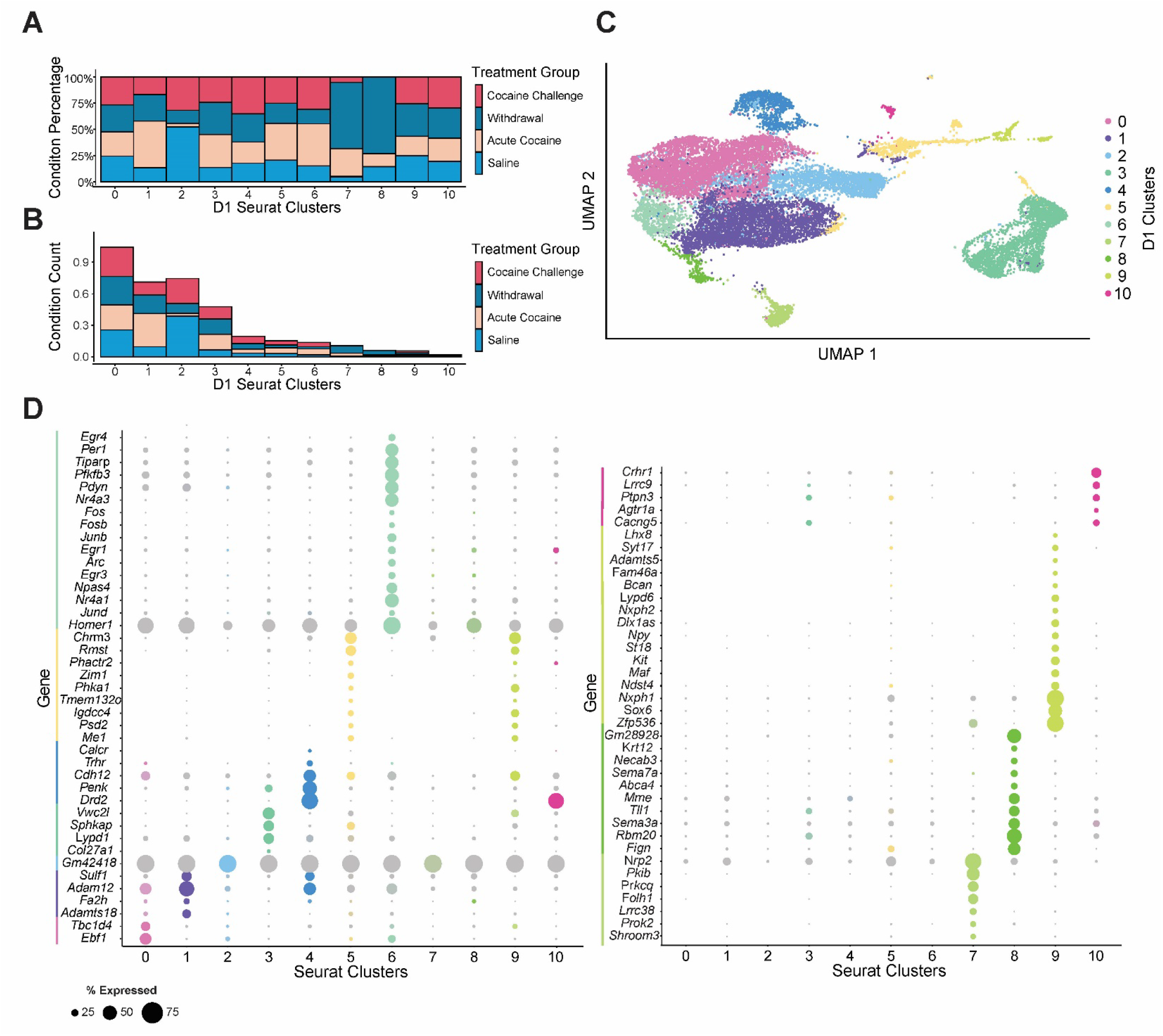
Transcriptomic analysis of D1 MSNs. A. Treatment condition percentage makeup of all D1 clusters. The bar chart shows the percentage of nuclei from each treatment condition (saline control, acute cocaine exposure, prolonged withdrawal, and cocaine rechallenge after prolonged withdrawal) within each of the 11 identified D1 MSN clusters, labeled from cluster 0 to cluster 10. **B.** Treatment condition total counts of the makeup of each D1 cluster. The bar chart quantifies the absolute number of nuclei from each treatment condition within each D1 MSN cluster, providing insight into the distribution of nuclei across different treatment groups. **C.** UMAP dimensionality reduction condensing all variable features into two dimensions for all D1 clusters. The UMAP plot illustrates the spatial arrangement and separation of the 11 D1 MSN clusters, reflecting the transcriptional heterogeneity within the D1 MSN population at baseline and in response to cocaine. **D.** Marker genes for D1 clusters. The dot plots are split into two sections for clusters 0-6 and 7-10. The size of each dot represents the percentage of nuclei within each cluster expressing the marker gene, while the color intensity indicates the average expression level of the marker gene within the cluster compared to all other clusters.

We further analyzed the quantitative distribution of nuclei from each treatment group across these transcriptionally distinct clusters within the D1 MSN population (Figure 2B), visualized in the UMAP plot following dimensionality reduction (Figure 2C). This visualization confirms the existence of unique clusters enriched under various treatment conditions. Notably, marker gene analysis identified specific genes that uniquely characterize each D1 cluster. For example, cluster D1_6 is distinctly marked by the expression of immediate early genes (IEGs) such as *Homer1*, *Fosb*, *Nr4a3*, and *Arc* (Figure 2D)^24,30,34,35^. The unique transcriptional identity of each D1 MSN cluster, highlighted by these marker genes, provides a comprehensive overview of the transcriptomic diversity within D1 MSNs and underscores the differential impact of cocaine treatment conditions on their gene expression profiles (Figure 2D).

### Transcriptomic Diversity of D2 MSNs

To further deepen our understanding of the transcriptomic diversity and the impact of cocaine treatment on MSNs, we conducted a comprehensive analysis of D2 MSNs, paralleling our approach with D1 MSNs. We examined the percentage composition of cells from each treatment condition across the nine identified D2 MSN clusters, revealing significant variations in the distribution of treatment conditions among these clusters (Figure 3A). For instance, cluster D2_2 exhibited a higher proportion of nuclei from the cocaine rechallenge group, suggesting long-term adaptations following prolonged withdrawal that may affect the immediate transcriptional response to cocaine upon re-exposure. This is further reflected by the relative distribution of nuclei across each D2 MSN cluster, where cluster D2_2 is predominantly composed of nuclei from mice subjected to a cocaine rechallenge (Figure 3B).

**Figure 3:**
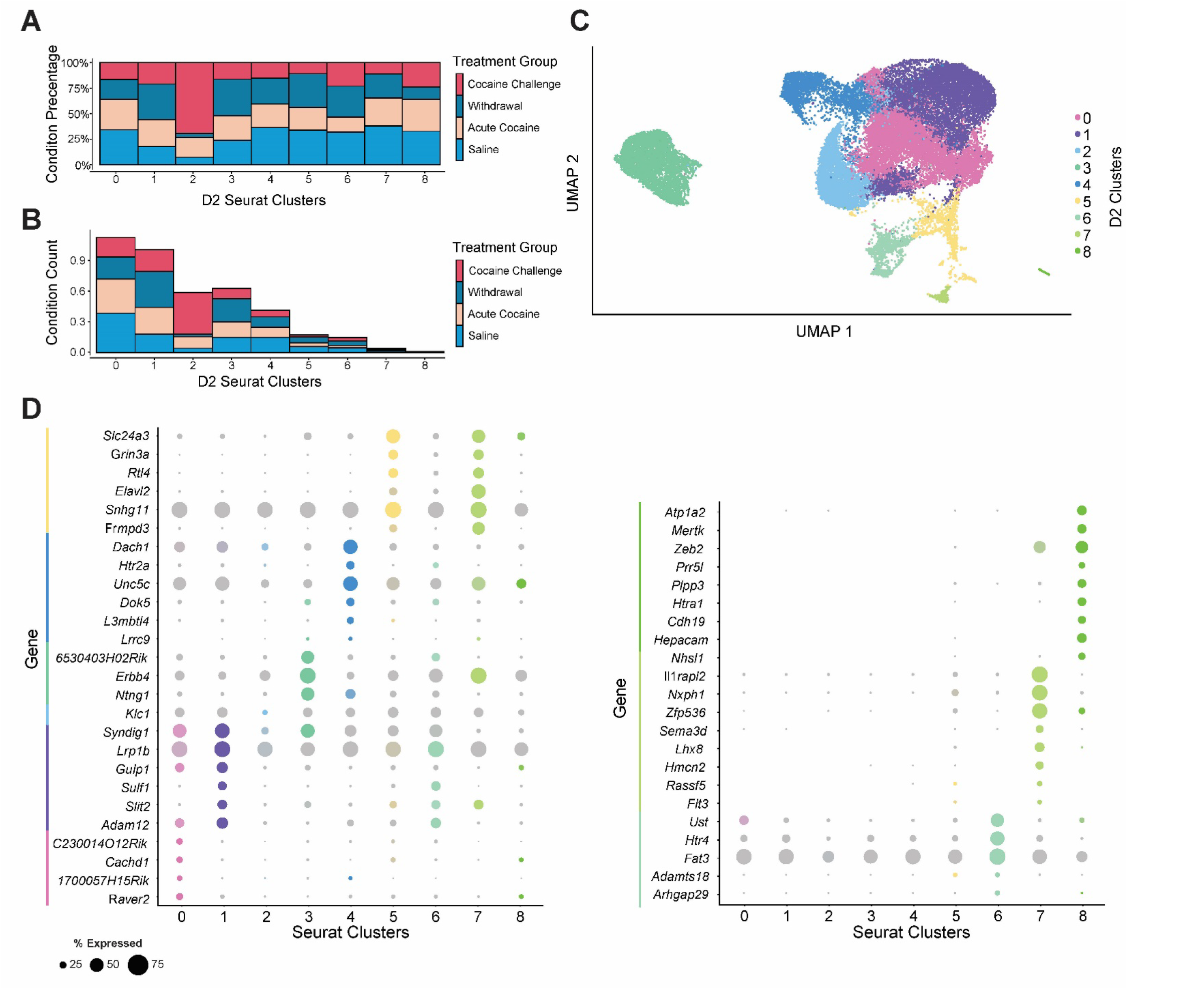
Transcriptomic analysis of D2 MSNs. A. Treatment condition percentage makeup of all D2 clusters. The bar chart shows the percentage of nuclei from each treatment condition (saline control, acute cocaine exposure, prolonged withdrawal, and cocaine rechallenge after prolonged withdrawal) within each of the 9 identified D2 MSN clusters, labeled from cluster 0 to cluster 8. **B.** Treatment condition total counts of the makeup of each D2 cluster. The bar chart quantifies the absolute number of nuclei from each treatment condition within each D2 MSN cluster, providing insight into the distribution of nuclei across different treatment groups. **C.** UMAP dimensionality reduction condensing all variable features into two dimensions for all D2 clusters. The UMAP plot illustrates the spatial arrangement and separation of the 9 D2 MSN clusters, reflecting the transcriptional heterogeneity within the D2 MSN population at baseline and in response to cocaine. **D.** Marker genes for D2 clusters. The dot plots are split into two sections for clusters 0-6 and 7-8. The size of each dot represents the percentage of nuclei within each cluster expressing the marker gene, while the color intensity indicates the average expression level of the marker gene within the cluster compared to all other clusters.

The distinct transcriptional profiles within the D2 MSN population were validated by identifying specific marker gene expression profiles for each D2 cluster (Figures 3C-D). Notably, these marker genes include well-characterized cocaine-responsive factors such as *Ntng1*, *Elavl2*, and *Grin3a*, which presumably denote specific transcriptional identities and functional roles of D2 MSNs, underscoring the heterogeneous nature of D2 MSNs and their differential transcriptional responses to cocaine exposure^6,36–38^.

### Unique Transcriptional Identities of a D1 MSN Cluster

To comprehensively profile the specific transcriptional dynamics within distinct D1 MSN subpopulations, we conducted an in-depth analysis of several clusters that exhibited discrete spatial separation or dominance of nuclei from key treatment groups. We began by examining cluster D1_3, which showed dramatic spatial separation from other D1 MSN clusters in the D1 UMAP and contributions from all three cocaine treatment groups. Analysis of cluster-specific marker genes revealed that this population was characterized by increased expression of key genes linked to synaptic plasticity. These include *Kirrel*, *Lypd1*, *Plcl1*, *Sphkap*, and *Vwc2l*, whose spatial distribution and expression levels across the D1 MSN population are illustrated in UMAP and violin plots (Figure 4A-B). Notably, *Kirrel* encodes a synaptic adhesion molecule critical for synapse formation that has been previously shown to be upregulated by cocaine.

**Figure 4:**
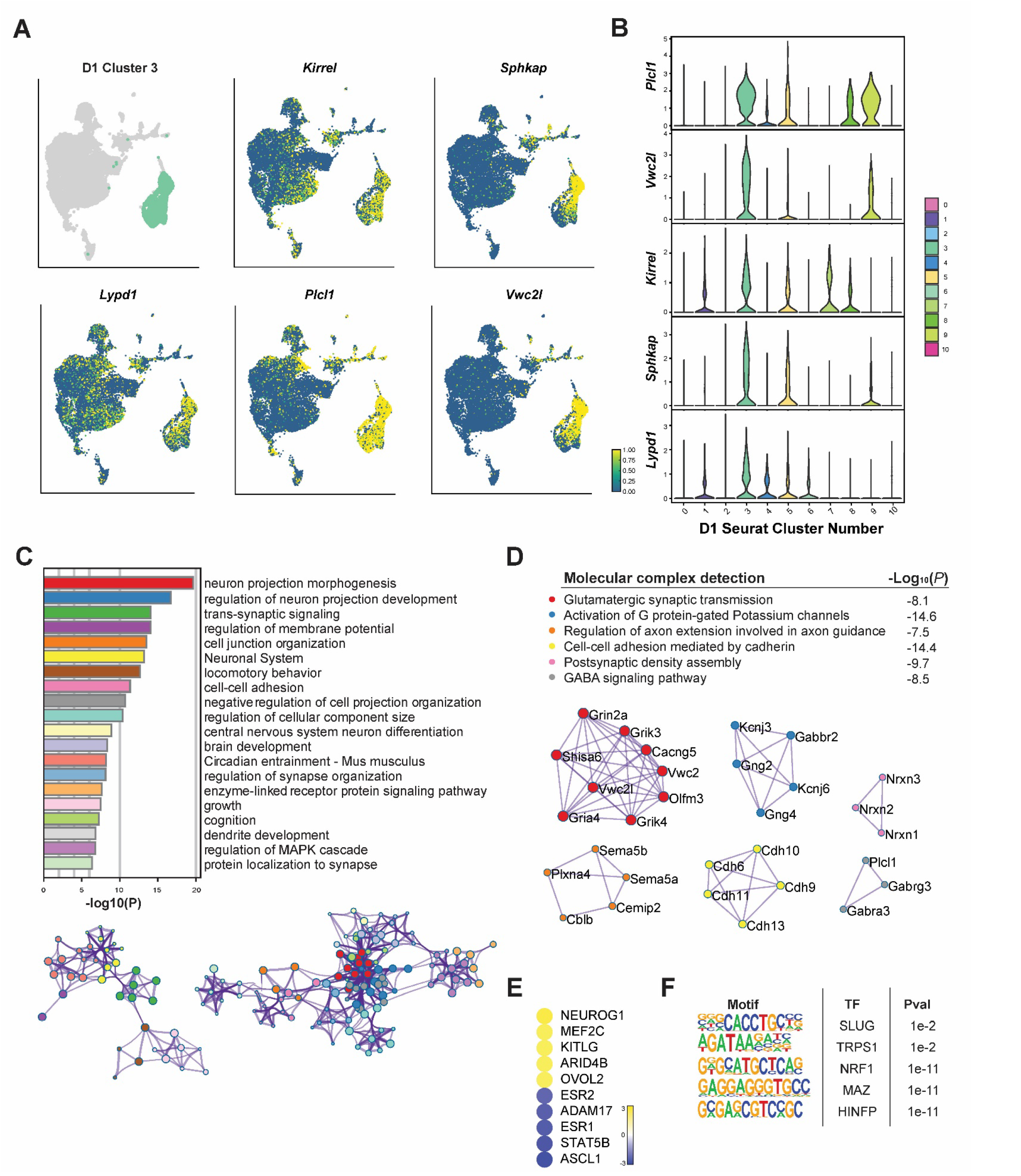
Detailed analysis of a transcriptionally distinct cluster of D1 MSNs. A. Expression of selected top marker genes in cluster D1_3 nuclei within the D1 MSN UMAP; this cluster is spatially very distinct from all other D1 clusters. The UMAP plot shows the spatial distribution and expression levels of selected top marker genes such as *Kirrel*, *Lypd1*, *Plcl1*, *Sphkap*, and *Vwc2l*. The intensity of color indicates the expression levels, illustrating the specific localization and transcriptional identity of cluster D1_3 within the D1 MSN population. **B.** Differential expression of top marker genes in cluster D1_3. The bar chart displays the -log10(P) values for the top differentially expressed marker genes, including *Kirrel*, *Lypd1*, *Plcl1*, *Sphkap*, and *Vwc2l*. **C.** GO term enrichment for cluster D1_3. The bar chart illustrates the -log10(P) values for enriched GO terms, including biological processes. **D.** Molecular complex detection on upregulated marker genes identified molecular mechanisms participating in synaptic transmission. **E.** IPA upstream regulator analysis performed on the significant marker genes identified key upstream regulators, such as *Nuerog1, Arid4b,* and *Ovol2*. This analysis provides insights into the regulatory networks governing this cluster. **F.** Enriched transcription factor motifs upstream of marker genes for cluster D1_3. Significant TF motifs include SLUG, NRF1, MAZ, and HINFP.

To provide additional insights into the functional roles of the upregulated marker genes that characterize D1_3, we performed Gene Ontology (GO) term enrichment analysis. These roles include biological processes central to neural plasticity, such as neuron projection morphogenesis, trans-synaptic signaling, regulation of the MAPK cascade, and dendrite development (Figure 4C). These enriched terms implicate the recruitment of key processes known to regulate synaptic plasticity and neuroadaptive responses to cocaine.

To elucidate the molecular mechanisms activated within the distinct population of D1_3 MSNs, we conducted molecular complex detection, identifying several complexes involved in synaptic signaling (Figure 4D). The most prominent complex detected is associated with glutamatergic synaptic transmission and includes the NMDA receptor subunit GRIN2A, which plays a critical role in synaptic strength modulation underlying addictive behaviors^39–41^. Similarly, cocaine has been previously shown to regulate the kainate receptor subunits GRIK3 and GRIK4, known for their roles in excitatory neurotransmission and linked to neural adaptations associated with addiction^42,43^. Another significant component of this complex is GRIA4, an AMPA receptor subunit, which is also impacted by cocaine and contributes to the synaptic changes observed in cocaine addiction^44–46^.

To further understand the regulatory networks governing the expression of these genes in cluster D1_3, we performed an upstream regulator analysis using Ingenuity Pathway Analysis (IPA). This analysis highlighted several key upstream regulators, including MEF2C, a transcription factor (TF) pivotal for synaptic plasticity and activity-dependent gene transcription critical for synaptic function. Cocaine has been shown to induce MEF2C in the rat striatum through activation of SIK1, resulting in synaptic structure and function alterations^47^. Conversely, the transcription factor ASCL1, involved in neurogenesis, is predicted to be inhibited in this population, consistent with cocaine exposure downregulating ASCL1^48^.

Finally, motif analysis identified TF motifs enriched upstream of marker genes upregulated in this distinct D1 MSN subpopulation. The analysis revealed that this transcriptional identity is regulated by factors such as SLUG, TRPS1, NRF1, MAZ, and HINFP. NRF1 is crucial for regulating mitochondrial function and oxidative stress responses, with cocaine shown to induce *Nrf1* in D1 MSNs while downregulating it in D2 MSNs^49–51^. MAZ, another TF regulating growth and differentiation genes, has also been linked to cocaine regulation^6,52^. Collectively, these upstream regulators and enriched TF motifs suggest complex regulatory networks contributing to the unique transcriptional identity of this D1 subpopulation.

### Cocaine-Responsive Immediate Early Gene Cluster of D1 MSNs

Based on its distinct transcriptomic profile, we identified cluster D1_6 as a relatively small subpopulation of D1 MSNs characterized by activity-regulated gene expression and enriched in nuclei from the acute cocaine and rechallenge conditions. This cluster’s top marker genes include well-known cocaine-responsive IEGs such as *Npas4*, *Arc*, *Fosb*, and *Nr4a1* (Figure 5A-B)^53–58^. For instance, *Arc* is highly responsive to neuronal activity and is induced by cocaine exposure, playing critical roles in cocaine-related synaptic plasticity and behavioral sensitization observed in addiction^53,59–61^.

**Figure 5:**
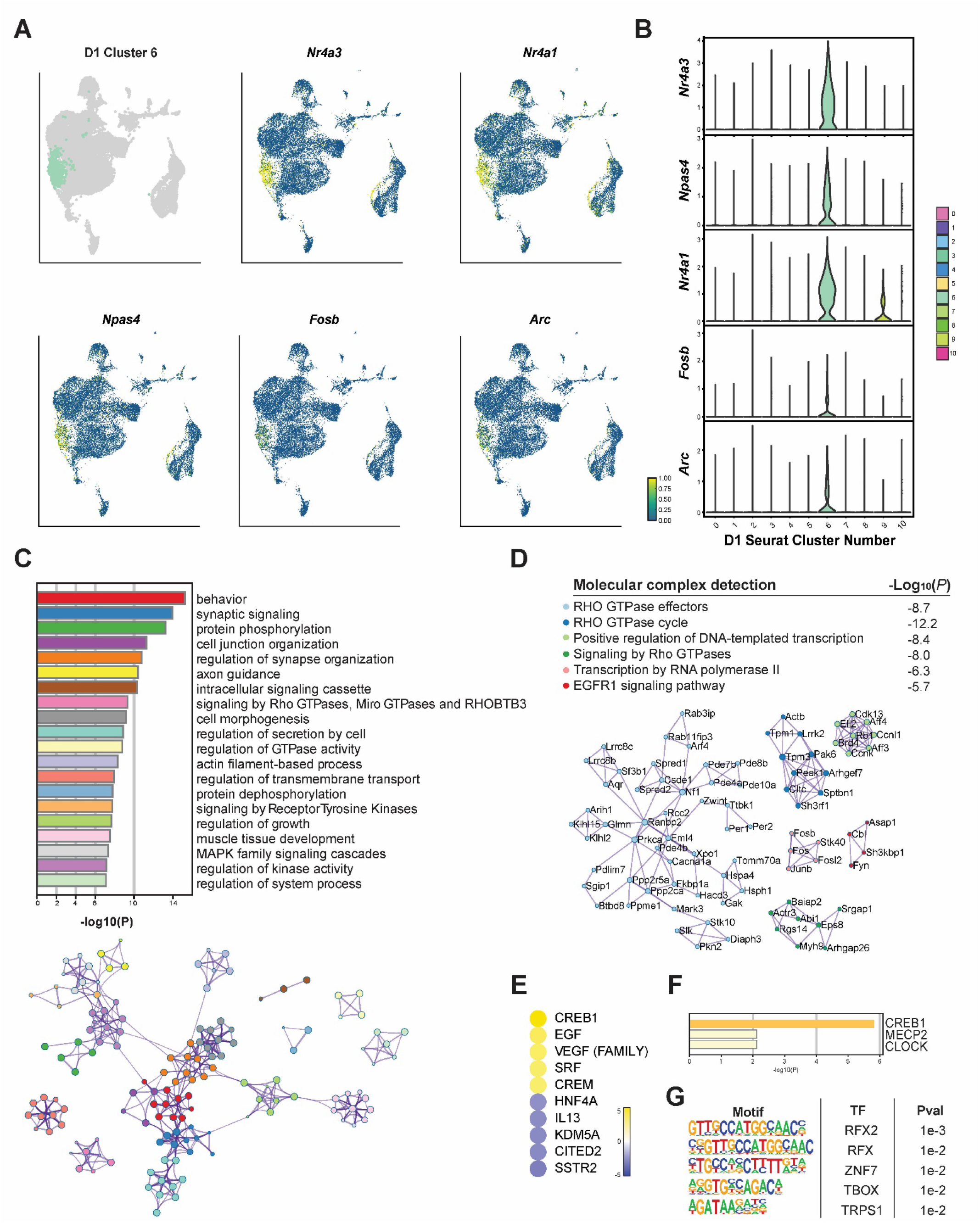
Detailed analysis of an IEG-enriched cluster of D1 MSNs. A. Expression of selected top marker genes in cluster D1_6 nuclei within the D1 MSN UMAP. The UMAP plot shows the spatial distribution and expression levels of selected top marker genes: *Nr4a1*, *Nr4a3*, *Arc*, *Fosb*, and *Npas4*. The intensity of color indicates the expression levels, illustrating the specific localization and transcriptional identity of this IEG cluster within the D1 MSN population. **B.** Differential expression of the top marker genes in cluster D1_6, marking them as key identifiers. **C.** GO term analysis for cluster D1_6, showing enrichment of biological processes including protein dephosphorylation, signaling by RHO GTPases, MAPK family signaling cascades, and axon guidance. **D.** Molecular complex detection of upregulated marker genes identified novel RHO GTPase effectors linked to cocaine action in the IEG cluster. **E.** IPA upstream regulator analysis highlights key upstream regulators, such as CREB1, EGF, VEGF, SRF, and CREM, implicating their regulation in cocaine-induced transcriptome dynamics. **F.** Transcriptional regulation predicted by TRRUST. **G.** Enriched TF motifs upstream of upregulated marker genes for cluster D1_6.

Similarly, neuronal activity rapidly induces *Fosb*, and its *ΔFosb* isoform in particular accumulates with repeated cocaine exposure and is crucial for long-term neural and behavioral adaptations associated with addiction^18,35^. Cocaine is known to epigenetically activate *Nr4a1*, which is integral to neuronal homeostasis and neuroprotection in response to hyper- excitation^57,62^.

GO network analysis of the upregulated marker genes in cluster D1_6 revealed involvement in biological processes central to activity-related plasticity, including synapse organization and Rho GTPase signaling (Figure 5C). Molecular complex detection further highlighted the recruitment of mechanisms controlling activity-dependent signaling and synaptic plasticity, identifying numerous RHO GTPase effectors (Figure 5D). Additionally, we found that key transcriptional regulators are active in this subpopulation, including BRD4, which interacts with acetylated histones to support the transcription of cocaine-responsive genes and is implicated in cocaine-induced reward^63,64^.

Upstream regulator analysis identified both canonical and novel cocaine-induced signaling pathways underlying the transcriptome dynamics in this subpopulation, including CREB1, EGF, HNF4A, CREM, and KDM5A (Figure 5E). Cocaine exposure activates CREB1 via the cAMP and CaMK signaling pathways, leading to its phosphorylation and activation, and the initiation of gene programs that control cocaine’s rewarding effects and contribute to addiction^6,65–67^.

Transcriptional regulation by CREB1 is further supported by TRRUST, a curated database of human and mouse transcriptional regulatory networks (Figure 5F). Similarly, CREM is a predicted activator that regulates plasticity-related gene expression in response to cAMP levels. Identification of HNF4A as an upstream regulator likely refers to the role of RXRa which we have shown recently is an important regulator in the NAc of cocaine’s transcriptional and behavioral actions^68^. In contrast, the epigenetic regulator KDM5A, a transcriptional co-repressor that demethylates the active chromatin mark H3K4me3^69^, is predicted to be less active in this D1 MSN cluster. Notably, epigenetic changes in both activating and repressive chromatin marks are thought to contribute to the long-term molecular and neural adaptations underlying addiction.

Our TF motif analysis further implicated the involvement of the RFX family of transcription factors, including RFX2 (Figure 5G). RFX2 is known to play a role in regulating genes involved in ciliary function and neuronal signaling^70,71^, suggesting its potential impact on the structural and functional adaptations in response to cocaine. Additionally, prominent zinc finger motifs, such as ZNF7, also emerged from this analysis. ZNF7 and other zinc finger proteins are involved in DNA binding and transcriptional regulation^72,73^, which are crucial for mediating the gene expression changes induced by cocaine.

These findings underscore the complex transcriptional regulatory networks that govern the immediate early gene cluster in D1 MSNs, implicating novel molecular factors in the synaptic plasticity linked to cocaine.

### Transcriptional Dynamics in a Cocaine Withdrawal-Associated D1 MSN Cluster

To elucidate the transcriptional dynamics within D1 MSNs linked to cocaine withdrawal, we focused on cluster D1_7, which displayed distinct spatial separation in the UMAP and was predominantly composed of nuclei from the withdrawal group. Detailed analysis of this cluster revealed a unique transcriptional profile characterized by several key marker genes involved in neuroadaptive processes, such as *Nrp2*, *Grik1*, *Stac*, *Rreb1*, and *Pkib* (Figure 6A-B). *Nrp2* is induced by repeated cocaine exposure and involved in axon guidance and synaptic transmission, potentially affecting neural circuit remodeling during withdrawal^74,75^. Another prominent marker regulated by cocaine is *Grik1*^42,76,77^, which encodes the glutamate receptor kainate-1 that participates in excitatory neurotransmission and is a target of topiramate, a medication used to reduce alcohol cravings. Relatedly, following chronic ethanol consumption, the PKA inhibitor, encoded by *Pkib*, has been found to be upregulated in the NAc^78^. STAC proteins, which regulate calcium channels and synaptic vesicle trafficking, though their involvement in cocaine-induced synaptic changes has been previously unknown, were also highlighted.

**Figure 6:**
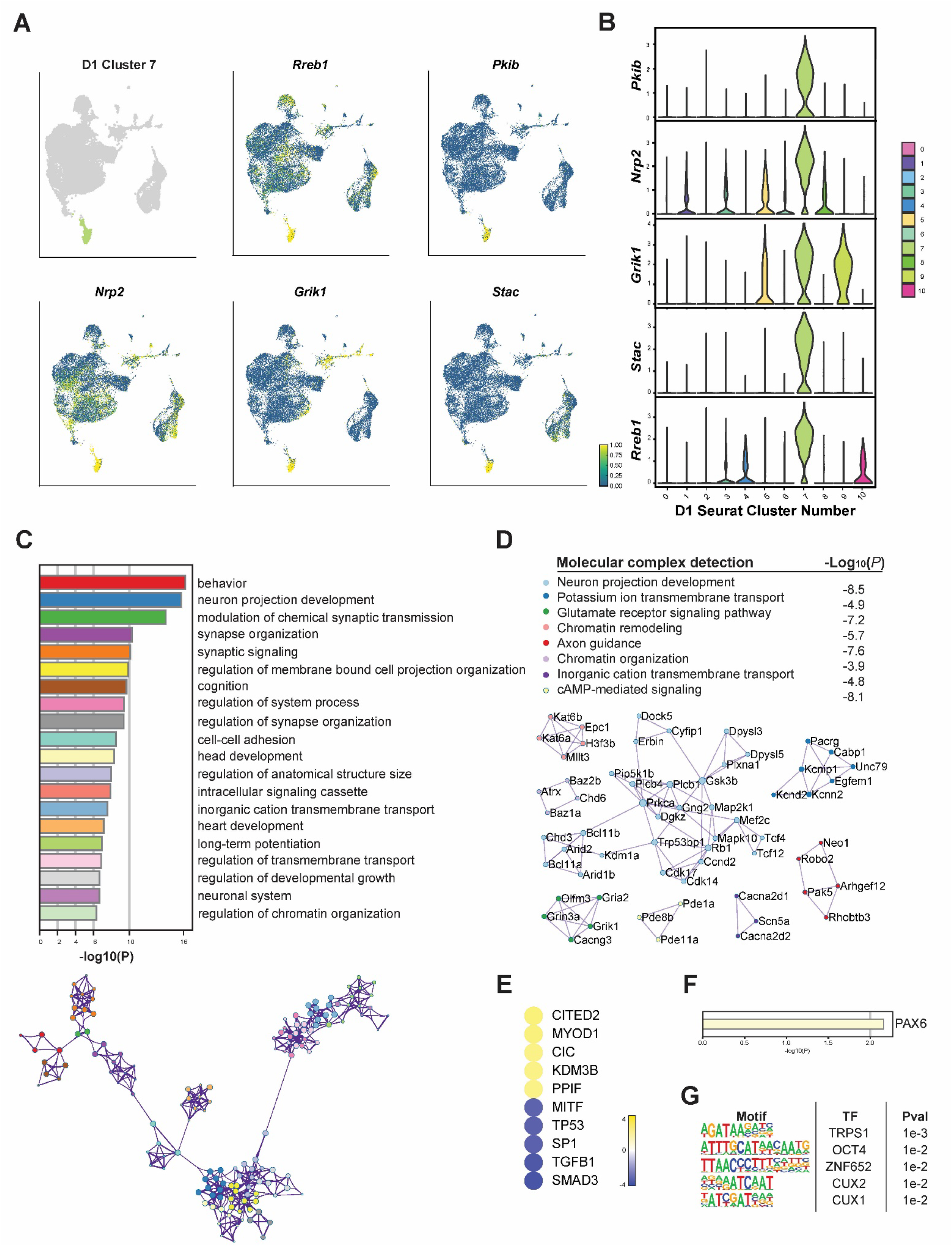
Detailed analysis of a withdrawal-associated cluster of D1 MSNs. A. Expression of selected top marker genes in cluster D1_7 nuclei within the D1 MSN UMAP, which shows the spatial distribution and expression levels of selected top marker genes: *Nrp2*, *Grik1*, *Stac*, *Rreb1*, and *Pkib*. **B.** Differential expression of the top marker genes in cluster D1_7. **C.** GO term enrichment for cluster D1_7, highlighting synaptic transmission, synaptic signaling, and regulation of chromatin organization as biological processes characterizing this cell population. **D.** Molecular complex detection of upregulated marker genes identified potassium and glutamate signaling, as well as chromatin remodeling, as key neuronal processes associated with withdrawal. **E.** IPA upstream regulator analysis performed on the significant marker genes for cluster D1_7 highlights CITED2, CIC, KDM3B, and PPIF as key regulatory activators governing the withdrawal-associated transcriptome. **F.** Transcriptional regulation predicted by TRRUST. **G.** Enriched TF motifs upstream of upregulated marker genes for cluster D1_6.

GO term enrichment analysis highlighted biological processes critical for the neuroadaptive changes associated with cocaine withdrawal, including modulation of synaptic transmission, synaptic signaling, and chromatin remodeling (Figure 6C). Molecular complex detection analysis further identified several key molecular complexes within cluster D1_7, particularly those involved in synaptic signaling via potassium channels as well as glutamate receptors, and in chromatin organization (Figure 6D). Prominent complexes included those associated with *Grin3a*, *Gria2*, *Kcnd2*, and *Robo2* (Figure 6D, green). The importance of GRIN3A in cocaine-induced synaptic plasticity and drug-seeking has been demonstrated using ex vivo patch-clamp recording, subcellular Ca^2+^ imaging, and transgenic mice^37,79,80^. Chronic cocaine has also been shown to alter AMPA receptor composition, including GRIA2, impacting synaptic strength and plasticity associated with addiction-related behaviors^8,44,46^.

Importantly, our analyses implicate novel epigenetic regulators in the transcriptome dynamics that characterize the withdrawal-associated D1 MSN cluster. These include the chromatin remodelers CHD3, CHD6, and ARID2, the histone demethylase KDM1A, and the histone acetyltransferases KAT6A and KAT6B. Chronic cocaine administration in mice has been found to induce histone hyperacetylation in the NAc at promoters of genes regulated by the NuRD complex containing CHD3^81^. In rats that had previously self-administered cocaine, KDM1A was found to be upregulated in the NAc^82^. Epigenetic changes mediated by these chromatin remodeling and histone-modifying enzymes are crucial in mediating the long-lasting differential transcriptional regulation observed during cocaine withdrawal.

Upstream regulator analysis identified several key regulators, including CITED2, CIC, KDM3B, and PPIF (Figure 6E). KDM3B is a histone lysine demethylase that specifically removes repressive histone H3K9me2/1 methylation^83^.

Expression levels of KDM3B and several other histone-modifying enzymes were previously found to be upregulated during early extinction training in rats that had previously self-administered cocaine^84^. This upregulation suggests a role for KDM3B in facilitating the transcriptional reprogramming necessary for behavioral extinction and synaptic plasticity following cocaine exposure. CITED2 is a transcriptional coactivator that interacts with CBP/p300, playing a significant role in the response to oxidative stress and modulating the activity of transcription factors involved in neuronal function^85^, such as SMAD and AP-2 proteins. CIC is another factor known to interact with transcriptional regulators to control gene programs linked to cell differentiation and stress responses^86^, which highlights the complex transcriptional regulation that occurs during cocaine withdrawal.

Finally, motif analysis revealed several enriched TF motifs upstream of the marker genes unique to cluster D1_7. These include motifs for TRPS1, ZNF652, and CUX2 (Figure 6F). CUX1 and CUX2 are homeodomain TFs that control dendritic branching, spine morphology, and synapse formation^87,88^. Together, these findings provide a comprehensive view of the transcriptional landscape within this withdrawal-associated subpopulation of D1 MSNs, highlighting the molecular mechanisms and regulatory networks that govern the unique adaptations observed during prolonged withdrawal from chronic cocaine. This analysis sets the stage for future investigations into targeted interventions that could mitigate the effects of cocaine use disorder by addressing the specific transcriptional and epigenetic changes identified in this study.

## DISCUSSION

Our study represents a significant advancement in understanding the transcriptomic changes induced by cocaine and prolonged abstinence across distinct subpopulations of MSNs in the NAc. By utilizing FANS coupled with snRNAseq, we achieved high-resolution analysis of D1 and D2 dopamine receptor-expressing neurons. This approach allowed for a detailed dissection of the complex and nuanced transcriptional responses specific to these neuronal subtypes under different treatment conditions, which is crucial for developing targeted therapeutic strategies for CUD.

Historically, discerning the specific cellular responses to cocaine within the diverse neuronal populations in the NAc has been challenging. Most prior studies of the transcriptional effects of chronic cocaine sequenced bulk NAc dissections and hence failed to capture differences that occur within specific neuronal subtypes^84^. At the other end of the spectrum, studies relying upon IEGs to identify drug-activated neuronal ensembles in NAc restricted their analysis to what we show is the small percentage of cells that display an IEG response. By contrast, by employing FANS of all D1 or all D2 MSNs coupled with snRNAseq, we isolated numerous subtypes of D1 and D2 MSNs that exhibit intricate changes in their gene expression profiles over time in response to various cocaine conditions. This approach thus identified many additional types of drug-activated ensembles – far beyond those detected previously based solely on IEG induction, work which highlights the dynamic and complex nature of transcriptional regulation in the NAc. Our findings also thereby extend previous research which indicated that D1 MSNs generally enhance reward-related behaviors, whereas D2 MSNs tend to suppress them; our findings provide a far more comprehensive and nuanced understanding by mapping how distinct subpopulations of D1 and D2 MSNs each respond over time to drug exposure, extended withdrawal, and rechallenge.

The use of the Seurat clustering algorithm unveiled significant heterogeneity within the D1 and D2 MSN populations, identifying distinct clusters that respond differently to cocaine exposure and withdrawal. For instance, clusters D1_5 and D1_6 showed a higher proportion of nuclei from the acute cocaine and cocaine rechallenge groups, while clusters D1_7 and D1_8 were more populated by nuclei from the prolonged withdrawal group. And cluster D2_2 was highly enriched in nuclei from the cocaine rechallenge group. These findings illustrate the complex transcriptional states that play different roles in the immediate and long-term responses to cocaine. Our results provide a comprehensive overview of the unique transcriptional identities of each cluster, offering insights into the specific molecular pathways and regulatory networks involved in cocaine addiction.

Moreover, the identification of specific transcriptional activators, such as CREM, RFX2, and ZNF7, through motif analysis of the immediate early gene cluster, informs the regulatory mechanisms underlying the persistent changes induced by cocaine. These transcription factors are implicated in a range of cellular functions, from DNA-binding to transcription regulation, suggesting their significant role in the long-term adaptations of MSNs to cocaine. Such insights are pivotal for developing interventions that target these specific molecular pathways, potentially offering more effective treatments for addiction tailored to the unique pathophysiology observed in individuals with CUD. Previous research has shown that transcription factors like CREB and ΔFOSB play critical roles in mediating the neuroplastic changes associated with cocaine addiction, further supporting our findings^18,34,67,89^. Our study also advances the understanding of transcriptional dynamics within MSN subpopulations following cocaine withdrawal. For example, we identified novel epigenetic regulators in a D1 cluster, including CHD3, KDM1A, and KAT6A, and implicate KDM3B, CITED2, and CUX2 in orchestrating the transcriptional reprogramming in response to repeated cocaine. These findings provide a comprehensive view of the transcriptional landscape and regulatory networks in this particular ensemble of D1 MSNs.

In conclusion, our study significantly advances the understanding of cocaine-induced molecular and transcriptional changes in the NAc by providing a detailed map of the transcriptional landscape in D1 and D2 MSNs during different phases of cocaine exposure, withdrawal, and rechallenge. By integrating advanced sorting and sequencing technologies, we have refined the understanding of how these neurons contribute to the addictive properties of cocaine and the potential pathways through which relapse may occur. This research underscores the value of high-resolution transcriptomic analysis in addiction studies and sets the stage for future investigations that could lead to novel therapeutic approaches for managing CUD. By offering new insights into the cellular and molecular mechanisms driving addiction and potential relapse, our findings pave the way for further research into targeted interventions for the treatment of cocaine and other psychostimulant use disorders.

## METHODS

### Animals

Male double-transgenic mice expressing D1- or D2-specific nuclear-tagged GFP were generated by crossing LSL-eGFP::L10a (IMSR_JAX:022367) mice with either D1-Cre (MGI:3836633) or D2-Cre (MGI:3836635) mice. All mice were bred in-house on a C57BL/6J background and were 8-20 weeks old at the beginning of experimental procedures. Mice were housed on a regular light-dark cycle with lights ON at 7:00 am, and food and water available ad libitum for the duration of the study. All experiments were conducted in accordance with the guidelines of the Institutional Animal Care and Use Committee (IACUC) at Mount Sinai (protocol number 08-0465).

### Cocaine treatment

Cocaine hydrochloride (from the National Institute on Drug Abuse) was dissolved in 0.9% saline and administered intraperitoneally at a dose of 20 mg/kg. This dose was chosen based on prior experiments (21). Mice received saline or cocaine each day for 10 days prior to withdrawal. As depicted in Figure 1A, after 30 days of homecage withdrawal, mice received challenge injections of saline or cocaine and were euthanized 1 hr later.

### Tissue processing and fluorescence-activated nuclei sorting (FANS)

Mice were euthanized by cervical dislocation, and bilateral NAc tissue was rapidly dissected on ice from 1-mm thick coronal sections using a 14G punch and flash frozen on dry ice. Nuclei were purified as previously described (21). Frozen NAc samples were first homogenized in lysis buffer (0.32 M sucrose, 5 mM CaCl2, 3 mM Mg(Ace)2, 0.1 mM EDTA, 10 mM Tris-HCl) using high-clearance followed by low-clearance glass douncers (Kimble Kontes). Homogenates were then passed through a 40um cell strainer (Pulriselect) into ultracentrifuge tubes (Beckman Coulter). A 5 ml high-sucrose cushion (1.8 M sucrose, 3 mM Mg(Ace)2, 1 mM DTT, 10 mM Tris-HCl) was then underlaid, and samples were centrifuged at 24,000 rpm at 4°C for 1h in a SW41Ti Swinging-Bucket Rotor (Beckman Coulter). After discarding the supernatant, nuclei pellets were resuspended in 500ul of ice-cold phosphate-buffered saline, and DAPI (1:5000) was added. All solutions contained inhibitors for RNAse (SUPERase-in, Invitrogen) and ribonuclease (RNasin Recombinant, Promega) at concentrations of 1:1000 (sucrose buffer) or 1:500 (lysis buffer and PBS). Nuclei were then immediately sorted using a BD FACS Aria II three-laser system with a 100um nozzle. Debris and doublets were first gated based on forward scatter and side scatter, nuclei were then gated as DAPI-positive events (Violet1-A laser), and GFP positive events were sorted directly into 1.5 ml Eppendorf tubes containing TriZol LS (Ambion) and flash frozen on dry ice. Samples ranged from 18,000-60,000 GFP+ nuclei collected for downstream analysis.

### Single-nucleus RNA sequencing

The snRNAseq data were imported as distinct 10X Genomics files for each respective sample. These files were subsequently integrated into a unified dataset while preserving the original sample identity, MSN type, and sample condition as metadata attributes. Cell Ranger filtered outputs were analyzed with Seurat v4.4.0^90^. The PercentageFeatureSet function was employed to calculate the mitochondrial gene percentage. Nuclei exceeding a 9% mitochondrial gene threshold were excluded to mitigate potential mitochondrial contamination that could influence marker analysis and dimensionality reduction outcomes. Data normalization was performed using Seurat’s LogNormalize method with a scale factor set at 10,000. A total of 2,000 variable features were selected using the "vst" selection method across the entire dataset. To further reduce potential contamination-induced alterations, data scaling was conducted with mitochondrial gene percentages regressed out. Principal Component Analysis (PCA) was initiated based on features derived from the "vst" method, and the Uniform Manifold Approximation and Projection (UMAP) computation employed 17 dimensions to ensure optimal cluster separation and distinct demarcation of D1 and D2 sorted populations within the integrated UMAP. The dataset was subsequently partitioned into D1 and D2 subsets for detailed analysis. UMAPs were recalibrated for these subsets, ensuring a coherent spread of clusters. The cell clustering pipeline was executed through Seurat, utilizing the Find Neighbors function with 17 dimensions, and determining clusters at a resolution of 0.25. This process yielded 11 clusters for D1 neurons and 9 for D2 neurons. These clusters were formulated to group cells with analogous gene expression profiles, facilitating the discernment of variations both inter-cluster and conditionally within a cluster. All datasets were archived as RDS files, accessible in supplementary materials. Intercluster analysis was conducted using Seurat’s FindMarkers function, comparing each cluster against all others within the same MSN type to identify gene expression disparities and potential cluster-specific genes. The cluster genes were also analyzed through BioMart v2.56.1 for basic gene information. Upregulated genes specific to each cluster were further analyzed through Metascape 3.5 for GO term enrichment analysis and molecular complex detection. All cluster marker genes were run through Homer for motif analysis to identify overrepresented transcription factor binding sequences.

### Metascape

Marker gene lists were subjected to functional enrichment analysis using Metascape 3.5 online platform^91^. These tools used advanced algorithms and curated databases to explore the biological processes and functional implications associated with the marker genes upregulated in select clusters. The analysis identified enriched GO terms, biological pathways, protein-protein interactions, and functional annotations, and included visually represented comprehensive networks and bar graphs, providing valuable insights into the molecular and cellular processes affected by cocaine exposure across D1 and D2 subpopulations.

## Data and materials availability

All data needed to evaluate the conclusions in the paper are present in the paper and/or the Supplementary Materials. All RNAseq data reported in this study will be deposited publicly in the Gene Expression Omnibus upon manuscript acceptance. Other supporting scripts/code used in this study are available from the corresponding author upon request.

## Acknowledgements

This work was supported by grants from the National Institute on Drug Abuse.

